# Divide and conquer: How avian “prefrontal” and hippocampal neurons process extinction learning in complementary ways

**DOI:** 10.1101/2025.03.21.644535

**Authors:** Celil Semih Sevincik, Julian Packheiser, José R. Donoso, Sen Cheng, Jonas Rose, Onur Güntürkün, Roland Pusch

## Abstract

Context-dependent extinction learning enables organisms to acquire an inhibition of responding to cues that no longer signal reward in specific environmental settings. Both the hippocampus and the prefrontal cortex play key but complementary roles in encoding extinction. To understand what drives the differential contributions of these two structures, we recorded single-unit responses from the pigeon hippocampus (HPC) and the ‘prefrontal’ nidopallium caudolaterale (NCL), while the animals were engaged in a repeated appetitive ABA extinction learning paradigm. HPC carried more information about experimental phases (acquisition (context A), extinction (context B), and renewal (context A)) than NCL. In addition, hippocampal neurons with mixed selectivity integrated all contextual stimuli and conditioned cues, thereby possibly enabling context integration into phase-dependent extinction events. The mixed selectivity pattern of NCL neurons differed: They encoded all relevant task parameters separately, thereby possibly enabling the weighing and deciding between response alternatives. Repeatedly testing the same animals over many months enabled us to reveal that the activity of NCL neurons increasingly became predictive of the number of responses that animals made during renewal. Thus, NCL neurons possibly assumed a meta-learning ability that enabled a behavioral adaption to the deeper task structure that always followed the sequence of acquisition→extinction→renewal. Our results make it likely that different processing strategies of hippocampus and ‘prefrontal’ NCL enable a differential encoding of experimental phase, context, decision-making, and meta-learning during extinction learning.

## Introduction

Animals constantly adjust their behavior to the ever-changing conditions of their environments. When, for example, an important food source is suddenly no longer available, animals stop visiting it and start looking for alternatives (Anselme & Güntürkün, 2019). The fading of a non-reinforced conditioned response is called extinction learning and can be emulated in experiments in which a previously acquired response to a conditioned stimulus is no longer rewarded such that animals extinguish their learned behavior and stop responding to the cue (Bouton et al., 2021). It is hypothesized that during extinction learning a secondary association is acquired that suppresses the acquired response to the conditioned stimulus (Quirk & Mueller, 2008; Bouton, 2004). Subsequently, the extinguished behavior can suddenly relapse when inhibition is reduced (Bouton et al., 2021; Milad & Quirk, 2002). These studies show that the originally acquired memory trace is not erased, even if the acquired behavior is completely abandoned. Although several variables contribute to the recovery of extinguished behavior, contextual cues are the most important ones and were thoroughly investigated in the literature under the term context-dependent extinction learning (Maren et al., 2013). The context-dependent recovery of a conditioned response, i.e. the renewal effect, is the most important relapse phenomenon under clinical conditions (Bouton, 2004). In laboratory studies, the initial learning of a conditioned response takes place in context A (e.g. white ambient light conditions). Subsequently, a new context B emerges (e.g. red ambient light conditions) in which conditioned responses are no longer rewarded. After a few trials, the animal gradually decreases its responses; the process of extinction learning has started. Subsequently, when the animal has entirely stopped responding to the conditioned cue and extinction learning has concluded, the context is changed back to A. Under these conditions, the animal vigorously starts responding to the conditioned cue again (Polack et al., 2013). For clinical therapy of anxiety disorders, renewal poses a serious issue, since the therapeutic intervention usually takes place in a different context (≈ context B) than the home of the patient (≈ context A) (Levy et al., 2022).

Accumulated evidence has shown that the neural substrates of context-dependent extinction learning are realized in a network comprising several brain regions involved in extinction and renewal (Kalisch et al., 2006; Phelps et al., 2004). Among these regions, the hippocampus (HPC) and the prefrontal cortex (PFC) have been identified as critical components. Previous lines of research reported that the HPC contributes to the integration of contextual cues into the extinction network since contextual cues serve to disambiguate conditions in which the CS+ may or may not be followed by a UCS (Fraser & Holland, 2019; Trask et al., 2017). Thus, the corresponding value of stimuli change in a context-dependent manner: While in acquisition context, a cue reliably predicts reward, this changes during extinction, where the changed context signals that no reward will follow the same cue. To produce adaptive behavior, context-dependent distinct neural representations of the same cue are required. Because hippocampal neurons also code for statistical regularities of feedback in decision-making tasks (Polti et al., 2022), the hippocampus may have the computational means to encode the context dependency of events in an extinction task.

The PFC plays another important role in extinction learning. The PFC is a key structure for executive functions and mediates adaptive behaviors when reward properties change (Dunsmoor et al., 2019). PFC neurons are thus strongly modulated during extinction recall (Milad & Quirk, 2002), while disruptions of prefrontal functions reduce the effect of extinction learning (Bukalo et al., 2015; Burgos-Robles et al., 2007; Morgan et al., 1993). Possibly, the PFC uses contextual information to adaptively guide behavior and initiate decisions, particularly in situations of ambiguity or conflict (Gonzalez & Fanselow, 2020). These flexible properties of the PFC are thought to be realized by neurons that integrate and evaluate all relevant task variables using multivariate response properties referred to as “mixed selectivity” (Rigotti et al., 2013). Besides individual contributions of the hippocampus and the PFC, the combined involvement of both regions was positively correlated with extinction memory recall (Milad et al., 2007), highlighting the network character of extinction learning.

But what enables the partly complementary contributions of hippocampus and PFC to extinction learning? Here, we assume that differences in the coding of behaviorally relevant task parameters drive complementary hippocampal and prefrontal roles during extinction. To test this hypothesis, we use pigeons (*Columba livia*) as model animals because they exhibit the same extinction-related learning mechanisms as humans and other mammals (Packheiser et al., 2019). In addition, pigeons learn very complex tasks that can be run over years in which they are willing to emit > 1,000 operant responses per daily session without a significant loss of motivation. Their moderate learning speed during our appetitive extinction learning paradigm allows a detailed trial-by-trial neurophysiological investigation of learning events (Packheiser et al., 2021). Moreover, neural and behavioral findings from experiments using avian species as subjects are often generalizable across species, including humans (Güntürkün et al., 2014; Kirschhock et al., 2021; Kirschhock & Nieder, 2022; Hahn et al., 2021; Güntürkün et al., 2024). The avian HPC is homologous to its mammalian counterpart and shares overlapping functions (Colombo & Broadbent, 2000; Payne et al., 2021; Applegate et al., 2023; Ben-Yishay et al., 2021). In contrast, the nidopallium caudolaterale (NCL) is not homologous to the mammalian PFC but constitutes an evolutionary analogous avian forebrain structure with highly similar neurochemical (Herold et al., 2011), dopaminoceptive (Durstewitz et al., 1998), connectional (Kröner & Güntürkün, 1999; Shanahan et al., 2013), and motor-control-related features (Steinemer et al., 2024). In addition, much behavioral and neurophysiological evidence supports the hypothesis that NCL and PFC share strong similarities at the neurobiological and functional level (Güntürkün, 1997; Diekamp et al., 2002; Hahn et al., 2021; Veit et al., 2014; Rose & Colombo, 2005). Consistent with our hypothesis, we find that NCL and HPC exhibit different patterns of mixed selectivity to behaviorally relevant task parameters, which might explain their complementary role in acquisition, extinction, and renewal.

## Methods

This paper reports new analyses of previous data collected by Packheiser et al. (2021) from NCL as well as novel data collected in this study from HPC specifically for this study. The experimental design and recording methods used in this study, and reported below, were matched as closely as possible to those in Packheiser et al. (2021) to make the two datasets comparable. For the analyses reported here, the data from both experimental studies were pooled and analyzed using the same methods.

### Subjects

Eleven naive homing pigeons (*Columba livia*) of mixed sex served in the experiments. They were housed in 30 x 30 x 45 cm wire-mesh cages separately. Water was provided ad libitum, whereas food access was restricted to experimental sessions. The animals were maintained at 80-90 % of their free-feeding body weight. Their normal weights ranged from 400 to 530 grams. If the animals performed poorly in the experimental sessions, they received additional food in their home cages to retain their bodyweights. During weekdays, they were fed with grains while they received a mixture of different kinds of corn on weekends. The colony room, where the aviaries were kept, was regulated by a 12-hr light-dark cycle, starting at 8 am. The subjects were treated in accordance with the German guidelines for the care and use of animals in science and all procedures were approved by a national ethics committee of the state of North Rhine-Westphalia, Germany. All experimental conduct agreed with the Directive 2010/63/EU of the European Parliament and of the Council of 22 September 2010 concerning the care and use of animals for experimental purposes.

### Apparatus

In the experiments, a custom-built 40 x 40 x 45 cm operant chamber was used. The boxes were illuminated from the ceiling with LED strips and had three horizontally aligned rectangular pecking keys (5 x 5 cm) on the rear side of the chamber. Below the middle key, a pellet feeder was located to provide food when required (http://www.jonasrose.net/open-labware/pellet-feeder/). The stimuli were presented via a small LCD monitor behind the transparent keys. The boxes were embedded into sound-attenuated wooden boxes (75 x 75 x 90 cm). Auditory feedback was delivered through external loudspeakers and white noise (∼60 dB) was present throughout all the experiments to prevent potential confounding variables. To run the experiments, custom-written scripts were used, and all the technical features of the boxes were controlled by the Biopsychology-Toolbox operating on MATLAB R2019a (Rose et al., 2008; Release 2019a, The MathWorks, Inc., Natick, Massachusetts, United States). All the statistical analyses were run in MATLAB.

### Behavioral Paradigm – Shaping procedure

Prior to the actual experiments, the animals underwent autoshaping and pre-training. During autoshaping, they first learned to associate the presentation of the initialization stimulus (an orange square) with food. The same procedure was repeated for the side keys. In both stages of the autoshaping, the animals had to provide correct responses at least 90 % of the time. Upon successful autoshaping, the animals proceeded to the pre-training with the following structure: First, the initialization stimulus was presented for 1 s. If the animals pecked during the presentation, one (out of two) sample stimulus (2.5 s) was presented. One stimulus was associated with a left key response, the other with a right key response. Animals learnt this association through trial-and-error. After the sample stimulus, they needed to peck one more time on the confirmation stimulus presented for up to one second (an orange square). In the last step, they needed to peck on one of the side keys to give a response. Given a correct response, we provided food via the pellet feeder. In case of wrong responses, we turned off the lights for two seconds, and no food was provided. If no response was given during these steps, the trial was aborted, and the inter-trial interval (4 s, ITI) commenced. In addition, we altered the lights in the chamber randomly (either to green or to red) at the beginning of each trial to familiarize the animals with the light changes that will occur in the actual experiments.

### Behavioral Paradigm – Extinction learning

To investigate extinction learning, we employed a classic ABA paradigm (Bouton, 2004) that was successfully adopted to pigeons (Packheiser et al., 2019). Every session consisted of three consecutive phases (Figure 1A), namely the acquisition phase (context A; gray background in figure 1A), the extinction phase (context B; red background in figure 1A), and the renewal test phase (context A; gray background in figure 1A). Each experimental session started with the acquisition phase where the animals had to learn the associations between the two novel stimuli presented during the sample phase and their respective assigned side keys (cf. Figure 1B). Two stimuli (control left, control right) were always the same in all the experiments. These stimuli kept the animals engaged in the task by ensuring a high performance rate in at least 50 % of the trials while learning about the left-right assignments of the novel stimuli, which were session unique (novel left, novel right). The side keys that the control items were associated with were counterbalanced between the animals. Control stimuli were rewarded in all phases (acquisition, extinction, or renewal), if a correct response was delivered. By contrast, for every session, the novel items were randomly assigned to be either the extinction stimulus or the non-extinction stimulus (Figure 1B). To proceed in the experiments, the animals had to perform correctly >85 % in a window of the past 100 trials for all of the four items (control and novel stimuli) during the acquisition phase. If not, the length of the phase was automatically prolonged until they fulfilled this criterion. To signal to the animals that the upcoming trial belongs to the context B, all the extinction trials were started with a light change (either to green or to red; Figure 1D) which remained until the end of the trial. Then, the context was switched back to the white again until the initialization of the next trial. Within each session, only one of the two colors was used. The most important point in this phase was that the animals had to extinguish their responses toward the extinction stimulus. Even though they gave (formerly) correct responses, they received neither reward nor punishment and they had to wait until the intertrial interval (ITI) started. Once their formerly correct responses toward the extinction stimulus and trials where the pigeons omitted to respond fell under 20 % in a window of the past 100 trials, the behavior was considered extinguished. However, the performance had to remain >80 % for the non-extinction novel stimulus and >75 % for the control stimuli. Once the extinction phase was terminated, the subsequent renewal test phase commenced. In this experimental phase, we tested the animals for renewal behavior under the same conditions as in the acquisition phase (context A). To prevent a confounding reacquisition of the extinction stimulus, we provided no feedback for responses that had been correct and rewarded during acquisition. Consequently, during the renewal test phase, no reward was delivered for the extinction stimulus regardless of the animals’ choice. There were no changes in the reward contingencies of the remaining stimuli.

**Figure 1.**
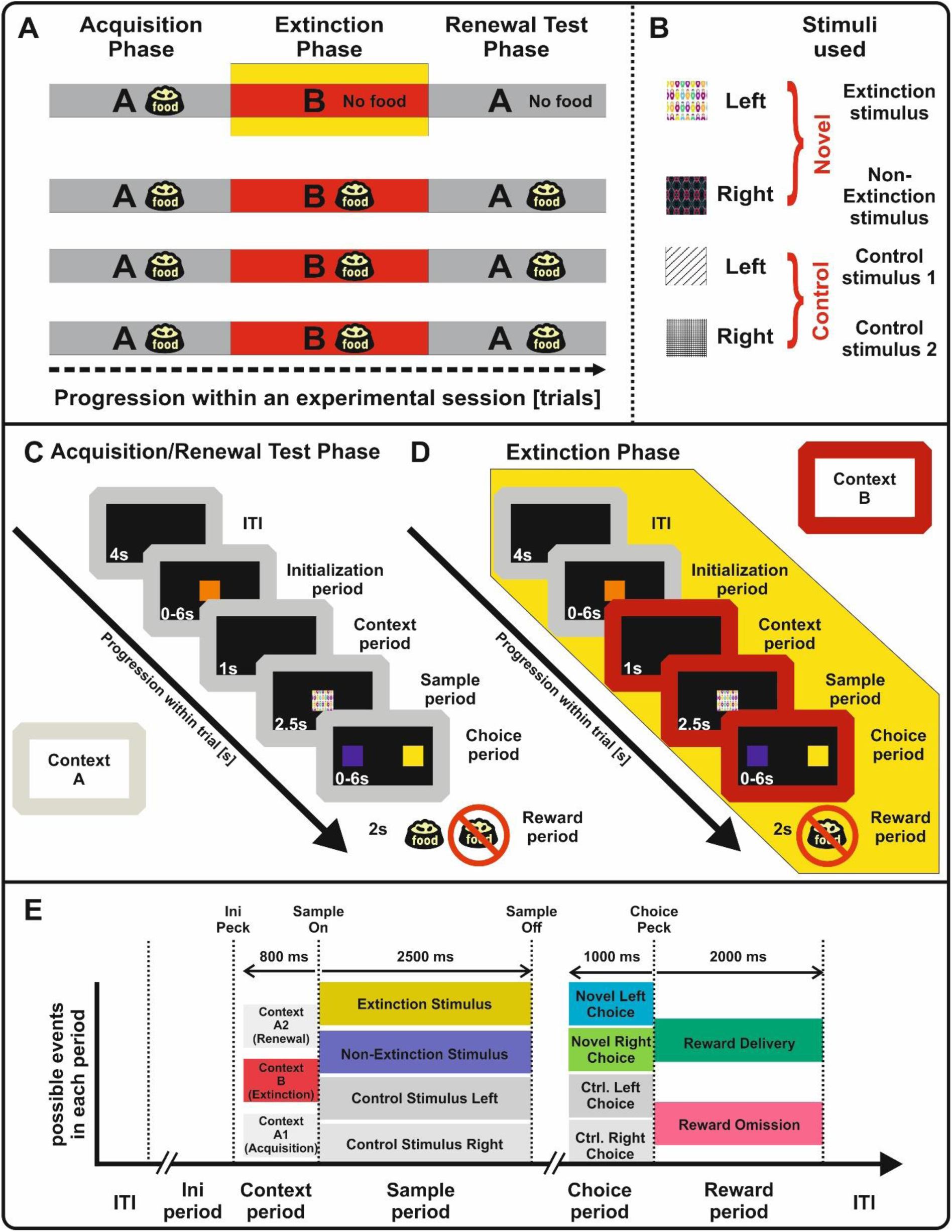
Illustration of the experimental paradigm**. (A)** Phases within a session. During the *acquisition phase* (context A) animals learned to associate left/right responses with two novel stimuli. After reaching a learning criterion, extinction started in context B (highlighted in yellow). One novel stimulus was randomly assigned to be extinguished (Extinction stimulus in panel B; trial structure in panel D). In the *renewal test phase,* context A was re-established, while pecking the extinction stimulus still did not yield reward (trial structure in Panel C). **(B)** The stimuli used in the experiment, their assignment to the side keys and the chosen extinction stimulus. **(C)** Trial structure and sequence of screen presentations during acquisition/renewal test phase. Please note that for the extinction stimulus, no reward is delivered regardless of the animals’ choice in the renewal test phase. **(D)** Trial structure and sequence of screen presentations during the extinction phase. An initialization peck on the center screen triggered a change in the light conditions to context B (red frame) 1 second before the stimulus presentation, which remained until the end of the trial. In extinction stimulus trials the outcome period remained void of any feedback regardless of the animal’s decision. **(E)** Parameter used in the linear regressions. Context was registered in three levels even differentiating between the first presentation of the context A in the acquisition phase and in the renewal test phase. Each stimulus was then coded separately. The decision responses were also differentiated whether they are directed to the left or to the right side as well as the stimulus. Lastly, we divided the reward parameters into two, namely the reward and no-reward conditions. Colors in each column correspond to the color-coding in figure 8.

### Electrophysiology

For our recordings, we used custom-made microdrives, each having one Teflon-coated silver (75 µm; Science Products, Hofheim, Germany) and sixteen formvar insulated nichrome wires (40 µm; California Fine Wire, Grover Beach, USA; Bilkey et al., 2003). Prior to the surgeries, the wires were gold-plated at the tips to acquire better signal quality (impedances ∼ 0.01 MΩ; Starosta, 2016). Before each session, we advanced the electrodes about 60 µm via a screw attached to the implant and waited for the unit responses to stabilize. Since each electrode on our implants could serve as a reference, we manually investigated all the channels to find the best signal quality. The extracellular potentials were first amplified 10x directly at the headstage and again 1,000x at the recording unit (EXT-16DX amplifier, NPI electronics, Tamm, Germany). The sampling rate was 22.08 kHz during all experiments. We filtered the raw signal within a window of 0.3-3 kHz during data acquisition. The data acquisition as well as offline sorting was performed on Spike2 software (Version: 8; Cambridge Electronic Design, Cambridge, UK). For the spike-sorting, we digitally band-pass filtered the raw traces (0.3-3 kHz) one more time and inspected their quality with custom-written MATLAB scripts (mlib toolbox, Maik Stüttgen, MATLAB central file exchange). To enter the final analyses, signals had to meet the following criteria to be considered as single-units: (1) the signal and the noise-band are separated by at least eight standard deviations which translates to signal-to-noise ratio of at least two. (2) Waveforms of the action potentials had to form a clear cluster in the principal component space. (3) The recordings had to be stable, having consistent spike amplitudes. (4) The recording had to be void of any disturbing movement artifacts e.g., discharges in close temporal relation to the pecking responses (Starosta et al., 2014). Since our analyses aimed to investigate the up or down regulation of the neural responses depending on the experimental phases, we additionally controlled for the stability of our recordings. To this end, we calculated the mean waveform (peak-to-through) amplitude for each neuron in each phase. Afterward, we compared the spike amplitudes within each phase on population level with Friedman’s test.

### Surgery

After the pre-training sessions, the pigeons were implanted with custom-made microdrives (Bilkey et al., 2003). For the initial anesthesia (Serir et al., 2024), a mixture of Ketamine (Ketavet, 100 mg/mL; Zoetis, Germany) and Xylazine (Rompun, 20 mg/mL; Bayer, Germany) was injected with a ratio of 9:1 (0,075 mL per 100 g bodyweight). Additionally, 0.05 ml of Buprenorphine (Buprenovet Multidose, 0.3 mg/ml; Bayer, Germany) was injected. After cutting the feathers on the head, the animals were placed on the stereotactic apparatus and provided with a constant flow of Isoflurane (Forane 100 %, Abbott GmbH & Co. KG, Wiesbaden, Germany). Once the animals showed no pain reflex, we incised the scalp and exposed the skull. First, 5 to 8 stainless metal screws were placed over the skull and then a small craniotomy was performed on the target locations (HPC: AP + 6 mm, ML + 0.5 mm; NCL: AP + 6.0 mm, ML ± 7.0 mm; Karten and Hodos, 1967). The histological track reconstruction is given in the supplementary figure S1 and shows the track of the implanted electrodes in the dorsomedial region of the hippocampal formation (Rook et al., 2023). After the dura removal, the electrode tips were inserted. Lastly, the craniotomy site was covered with Vaseline, and the whole implant was stabilized with dental cement around the metal screws. For three days following the surgery, the animals were supplied with analgesics (Rimadyl, 50 mL/mL Carprofen; Zoetis, Germany) and allowed to recover for at least a week.

### Behavioral data analysis

To keep the findings between the datasets as comparable as possible, we adopted the methodology introduced by Packheiser et al. (2019). We first computed the percentage of the correct conditioned responses for each stimulus and phase for each session. Since each session varied in duration depending on the animals’ performance, we standardized them by dividing each experimental phase (the acquisition phase, the extinction phase, and the renewal test phase) into 6 blocks and then calculated the average for each block. Afterward, we probed the extinction of the conditioned behavior for the extinction stimulus by comparing the last block of the acquisition phase with the first block of the extinction phase with a paired sample t-test. In a similar vein, we contrasted the last block of the extinction phase to the first block of the renewal phase to test the existence of renewal behavior. We exercised the same procedure for the remaining blocks of the renewal test phase to probe the time span of the renewal behavior with successive paired sample t-tests.

### Neural data analysis

Because each neuron has a distinct baseline firing rate (FR) and each session differed in terms of the number of trials, we normalized our data to eliminate any bias that may have stemmed from these factors. Therefore, we chunked the neural activity into 100 ms bins and estimated the mean and standard deviation of FRs across trials during the ITI of each session for each particular stimulus. Then, we normalized the raw neural responses in each trial via z-transformation. The z-transformed FRs were used for all subsequent statistical analyses. The spike density functions were then plotted after smoothing to provide a descriptive view of the neural responses on a population level.

In our experiment, three main factors were manipulated:

1. The *phase* of the extinction learning process: The first phase was the acquisition phase in which animals learned to associate novel stimuli with a corresponding behavioral choice. The subsequent extinction phase followed, in which one of the novel stimuli was no longer rewarded in a different context. Finally, there was the context-dependent renewal test phase in which the contextual cues of the acquisition phase re-appeared without altering the reward contingencies (Figure 1A).
2. The *stimulus* of the experiment: We employed two familiar stimuli that were presented in every session and served as controls. In addition, two novel stimuli were introduced that had to be learned within the acquisition phase of the respective session (Figure 1B).
3. The *periods* within trials: In the context period, the contextual cue was present exclusively. In the subsequent sample presentation period, the stimulus appeared on the center response key. During the choice period, animals had to decide to peck to the left or right-side key. Finally, the reward period followed, in which either food, punishment, or no feedback was delivered (Figure 1C/D).

To compare the effects of the manipulated factors, a 3-way ANOVA on the z-transformed neural responses was conducted for each individual neuron. This analysis allows to investigate the main effect of each of the abovementioned factors. In addition, the co-occurrence of neuronal firing in response to two factors can be found as two-way interaction effects (e.g., the extinction stimulus is only responded to in the extinction phase). Even more complex three-way interactions (e.g., the extinction stimulus is only responded to in the extinction phase during the reward period where the expected food is omitted) can be investigated. We continued our analysis with effect sizes due to the following three reasons: (1) *p*-values are susceptible to various factors (such as number of trials and FR) and do not quantify the strength of a putative statistical significance. (2) They allow us to decode the differential contribution of each factor and interaction for each neuron. (3) They provide a common ground for the comparison of different neurons. Therefore, following each 3-way ANOVA we computed the proportion of the explained variance for each of the three main factors (*phase, stimulus, periods within trials*) as well as their interactions (four in total: phase*stimulus, phase*period, stimulus*period, 3-way interaction). More specifically, we calculated the corresponding ω^2^ values based on the sum of squares of the factors and the mean squares of the within-group (error) variance (Equation 1).

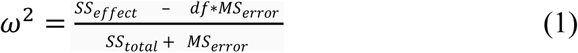

We preferred ω^2^ due to our multifactorial experimental design and its standing as a less biased measure of effect sizes (Olejnik & Algina, 2003). Consequently, we were able to compare the statistics of the different ANOVAs belonging to different neurons with each other. Hereafter, we will refer to the proportion of variance explained by each factor or interaction as *information* carried by that factor or interaction throughout the remaining text.

Once we quantified the information carried by each factor or interaction for each neuron, we could investigate the possible differences between both structures. To analyze the effect of the region (HPC vs. NCL) and the type of information (seven in total: phase, stimulus, periods within a trial, and all the interactions) on the amount of information carried, we performed a two-way mixed ANOVA (Figure 4).

### Multiple linear regressions

Although the 3-way ANOVA allows to investigate multiple combinations of activities as two-way or three-way interactions, this type of analysis fails to disentangle which parameters paired up dynamically. To investigate the manipulated factors in more detail, we computed multiple linear regressions with 13 distinct task parameters as predictors and raw spike counts as dependent variables. The parameters were divided into context, stimulus, decision, and reward parameters (cf. Figure 1E). This analysis allows us to disentangle, which task parameters altered the cells’ responses at a single trial resolution. In addition, the co-occurrence of multiple predictors can be investigated, resulting in the precise temporal response dynamics during extinction learning.

Context parameters were divided by experimental phase. During the acquisition phase, context A1 was measured during the last 800 ms of the 1-s-long context period of the experiment. The first 200 ms were excluded to remove potential residual activity from trial initialization. The same calculation was performed for context B during the extinction phase and context A2 during the renewal test phase. For stimulus parameters, we measured the number of spikes to each of the four experimental stimuli during the sample period of 2.5 seconds, namely the two familiar control stimuli and the two novel stimuli depending on whether they required a left or right choice. The same was true for decision parameters, but decision-related spiking was measured 1000 ms before the choice was made, i.e., during the choice period. This time window was chosen since numerous studies have demonstrated premotor activity patterns in the NCL before the chosen behavior was executed (Starosta et al., 2014; Lengersdorf et al., 2014a; Veit et al., 2014). Finally, reward parameters were subdivided into delivery and omission.

To determine the significance of the predictors, we permuted the raw spike counts randomly across the trial. These permutations were performed 35.360 times for the NCL cells as 1/35.360 is the likelihood of a predictor becoming significant at chance level with a Bonferroni corrected threshold (alpha = 5 % divided by 13 (predictors) * 136 (recorded cells). We used the same threshold for the HPC data to make the data analysis as comparable as possible between the two datasets. We considered a predictor significant if the empirical β-value resulting from the multiple linear regression exceeded all 35.360 β-values from the permuted data sets. All significant β-values were categorically transformed into +1 if the β-value was positive and -1 if the β-value was negative to reflect whether individual neurons increased or decreased their firing rate significantly for each predictor to obtain a significance matrix.

We then generated a graph in which each connection between two predictors indicated a significant coupling between those predictors. To obtain such a graph, we first generated an adjacency-count matrix in which each entry contained the number of cells that responded to any two predictors specified by a row and a column. The diagonal of such a matrix indicates the number of cells responding to any given predictor. To assess the significance of each entry in the adjacency-count matrix, we estimated the probability of obtaining the respective count under the null hypothesis that responses are randomly distributed across cells. To generate a distribution for the null hypothesis, we randomly permuted the entries of each row of the design matrix and re-ran the analysis. In this manner, we generated a random pattern of co-responses while conserving the number of cells responding to any given predictor. By repeating this procedure 10,000 times, we obtained a distribution of surrogate counts for each entry of the adjacency-count matrix. We considered an entry significant if its respective count was above the 99^th^ or 99.9^th^ percentile of the surrogate distribution. The final adjacency matrix describing the connectivity between predictors corresponds to the significant entries of the adjacency-count (see Supplementary Figure S2).

## Results

### Behavioral results

Previous research showed that that both HPC and prefrontal areas contribute to extinction learning (Lengersdorf et al., 2014b; Packheiser et al., 2021). To gain further insights into this process, we recorded single-unit responses from the HPC and the NCL while the animals were engaged in an identical appetitive extinction learning task. Comparisons and detailed analysis of such recordings can only be meaningful if the animals in both experiments exhibit the same behavior. Therefore, we started our analysis by comparing the behavioral sessions from both datasets. The depiction of the correct choices for the different stimuli in different phases can be found in figure 2. Here we provide the results stemming from the HPC dataset and present them together with the behavioral results from the NCL dataset (Packheiser et al., 2021; Donoso et al., 2021).

**Figure 2.**
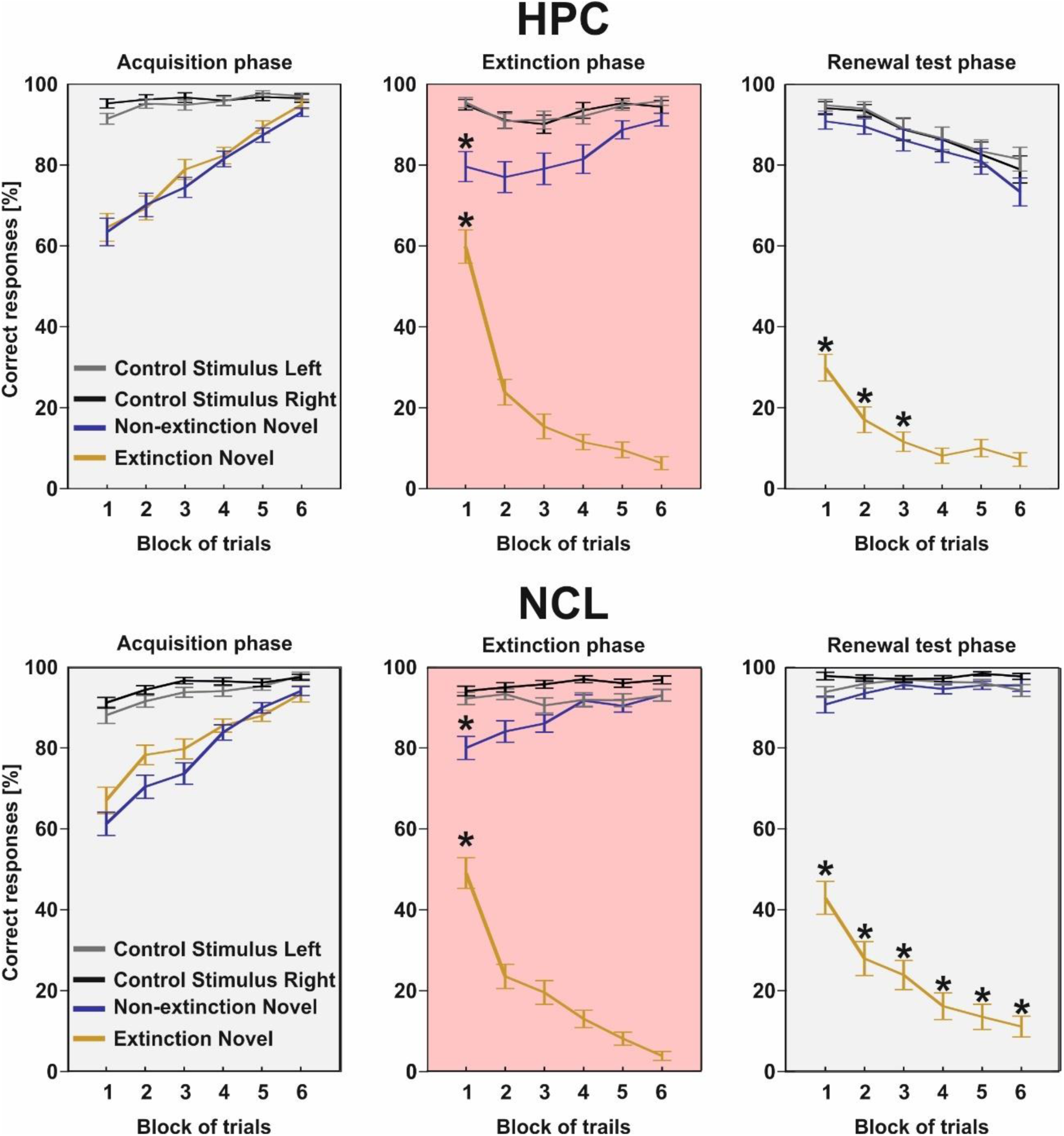
The depiction of the behavioral results from the HPC and NCL datasets. The leftmost panels show that while the control stimuli (Left Familiar, Right Familiar) were associated with the correct choices successfully from the very beginning of the acquisition phase, the pigeons learned the necessary associations for the novel items (extinction and non-extinction stimuli) over time. The middle panels depict the number of conditioned responses for the stimuli during the extinction phase. Most importantly it reflects the decline in the responses to the extinction stimulus, showing the extinction of the acquired behavior. The rightmost panels show the relapse of the conditioned behavior in the renewal phase. Asterisks mark where the number of conditioned responses differed significantly from the last block of the previous phase. The error bars represent the standard errors of the mean.

Since each session differed in duration depending on the animal’s performance, we divided each phase into 6 blocks. To investigate if and how extinction learning took place, we started our analysis by comparing the last block of the acquisition phase to the first block of the extinction phase. Already in the first block of the extinction phase, we found a significantly lower rate of conditioned responses to the extinction stimulus (paired t-test, *t*_(68)_ = 8.48, *p* < 0.001). Besides the responses to the extinction stimulus, we also found slightly fewer conditioned responses for the non-extinction stimulus (paired t-test, *t*_(68)_ = 3.74 *p* < 0.001). However, the animals reached the same success level as in the acquisition phase in the last block (paired t-test, *t*_(68)_ = 1.01, *p* = 0.32). This shows that the pigeons generalized their knowledge of both novel stimuli at the very beginning of the extinction phase and differentiated between them as they progressed in the experiment.

To test for renewal, the relapse of the conditioned responses after re-establishing the acquisition context A, we compared the last block of the extinction phase to the first block of the renewal test phase. We found significantly more conditioned responses for the first block of the renewal test phase, clearly showing the relapse of the extinguished behavior (paired t-test, *t*_(68)_ = -7.12, *p* < 0.001). The renewal effect was significant and could be detected in the first three blocks of the renewal test phase. Thereafter, no difference in the number of conditioned responses compared to the last block of the extinction phase could be detected due to the continuous omission of the expected reinforcer and the resulting re-extinction.

In summary, the pigeons implanted in the HPC showed identical behavioral patterns during the experiments and thus replicated the behavioral results of the pigeons that were implanted in the NCL (Packheiser et al., 2021).

### Electrophysiology

We then proceeded to analyze our electrophysiological recordings. For the current study, we recorded 98 HPC single-unit responses. The NCL dataset (n_NCL_ = 136) stems from Packheiser et al. (2021) and was analyzed in an earlier study with respect to prediction error-associated signals. Both the new and the previously published NCL-dataset were jointly analyzed in a novel way to investigate the real-time neural correlates of context-dependent extinction learning. The results have not been published elsewhere.

We first standardized the raw firing rates of both datasets via a z-transformation. Thus, we calculated the mean firing rates over trials within the ITI for each stimulus and subtracted it from the mean firing rates of each trial. These differences were then divided by the respective standard deviations. To have a population-level description of the neural responses, we plotted the normalized spike density functions of each neuron for each period in the sessions. The color-coded traces of the normalized neural activity in response to the extinction stimulus (Figure 3) indicate that the HPC shows less temporal specificity in its response profile as compared to the NCL. Furthermore, the HPC seems to be preferentially engaged during the context and stimulus presentation periods during the extinction and renewal test phase (Figure 3, lowermost row). Please also refer to the supplementary figure 3.

**Figure 3.**
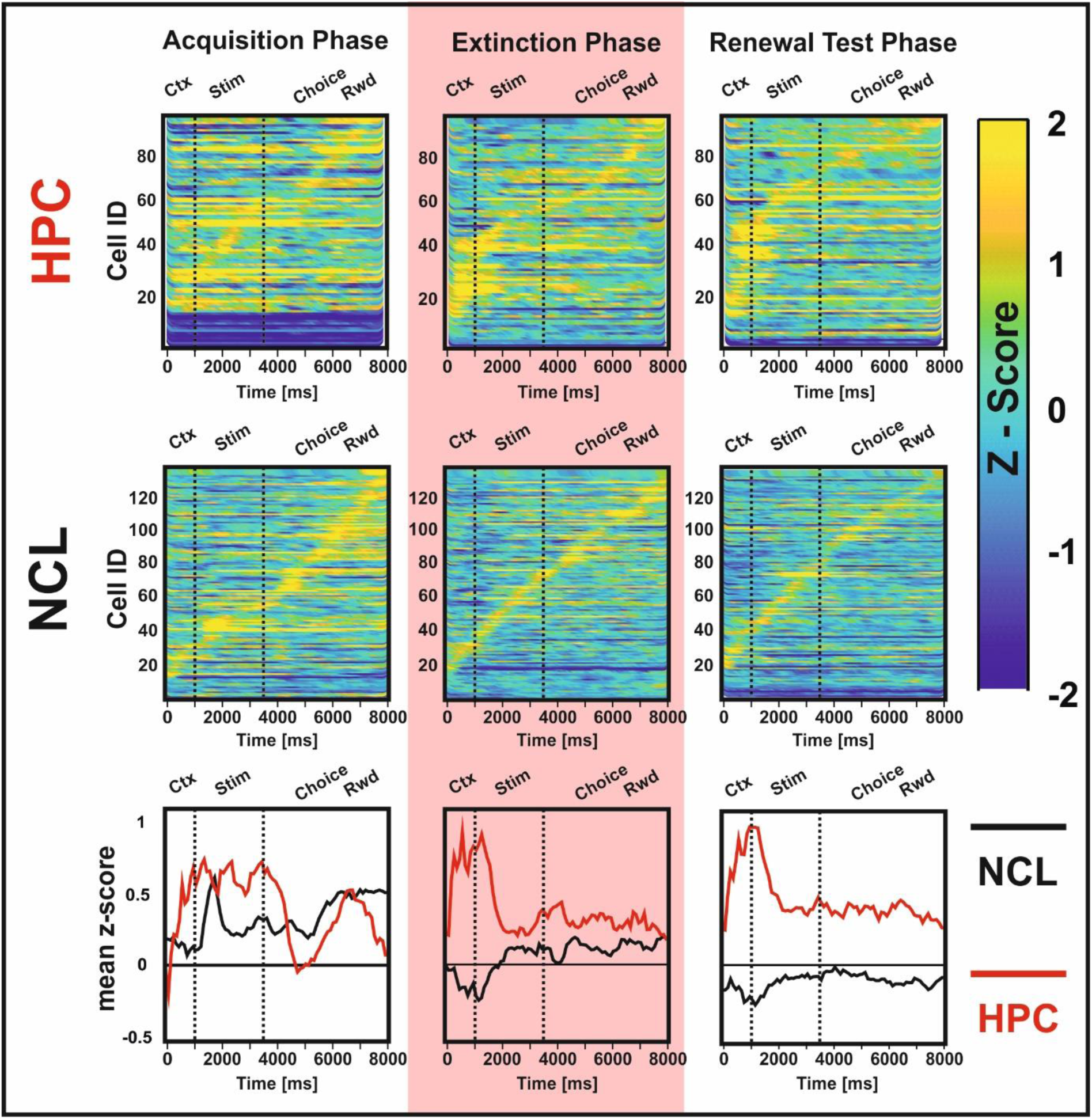
Depiction of population-level neural responses for the extinction stimulus for both datasets. The raw responses were first pooled together in bins of 100 ms and then standardized via z-transformation using the mean and standard deviation of the trials throughout the whole ITI. The spike density functions were then smoothed and sorted by the latency to the maximum value. The first row shows responses from all the HPC neurons, while the second row shows the same for the NCL. Each row within each panel corresponds to a single unit. The red background highlights the extinction phase. The third row depicts the mean z-score for each bin for the HPC (red traces) and the NCL (black traces). Overall, the NCL neurons preferred specific periods during the experiment, while the HPC neurons had a broader response profile. For the detailed population responses of both datasets toward the remaining stimuli please refer to supplementary figures 4 and 5.

To compare the response profiles between the two brain regions, we investigated the impact of our experimental manipulations on a single unit level. In our experiments, there were three experimental *phases*, namely the acquisition phase (context A) where animals acquired the association between each stimulus and the corresponding key; the extinction phase (context B) where the extinction stimulus was not rewarded anymore and therefore the conditioned responses were extinguished; and finally, the renewal test phase (context A again) where the animals were tested for the relapse of the previously extinguished behavior. In addition, we used two familiar control stimuli and two novel stimuli which directed the pigeons either to peck on the left key or on the right key to potentially earn a reward. Lastly, the paradigm comprised different experimental *periods* within each trial: exclusive presentation of the contextual cue, the stimulus presentation, the choice period, and finally the outcome period featuring reward, punishment, or no feedback (cf. Figure 1 C, D & E). To assess the effect of these factors on the z-transformed FRs, we conducted a three-way ANOVA on the aforementioned factors (phase x stimulus x period in a trial) for each neuron individually (Figure 4).

**Figure 4.**
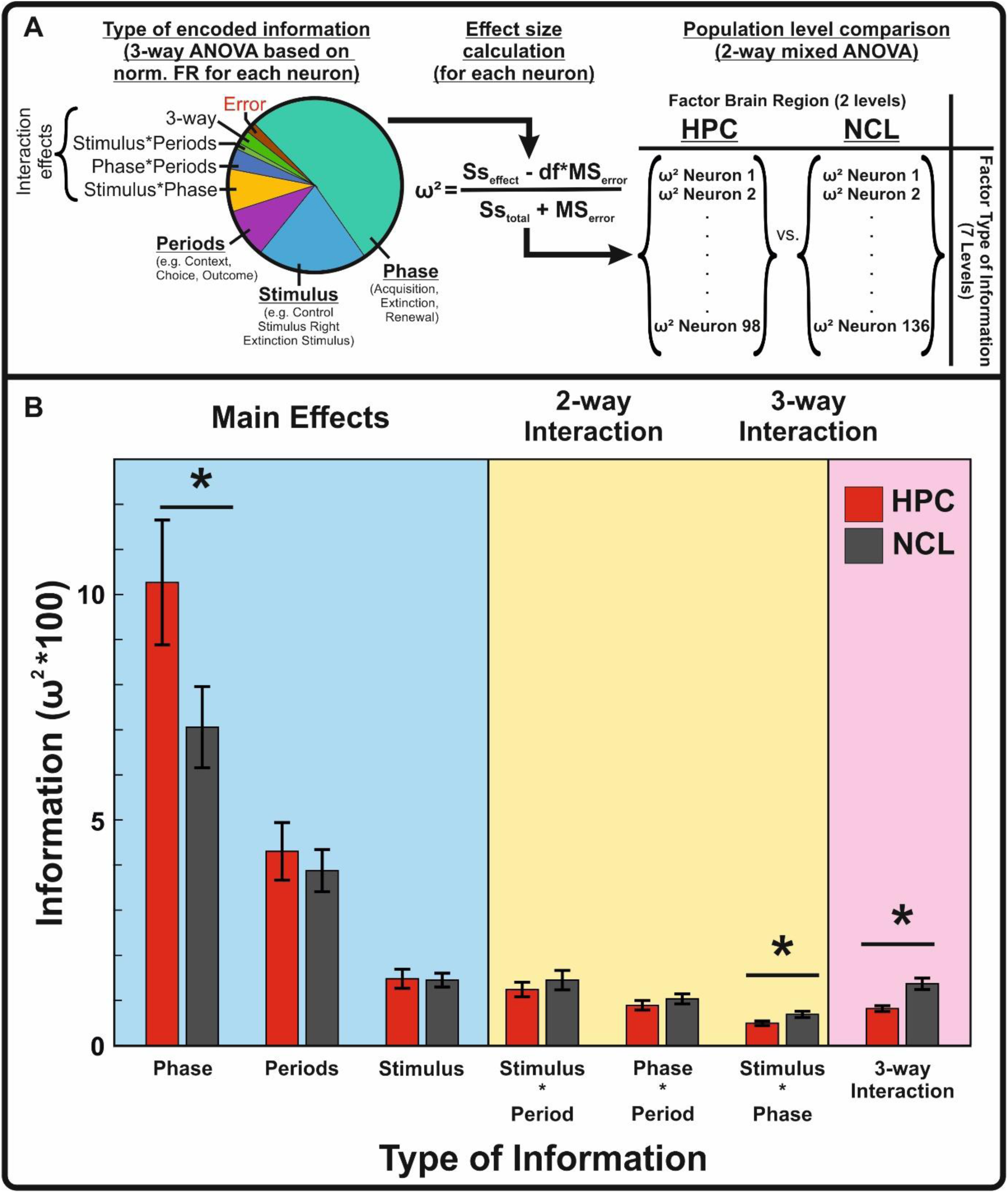
Depiction of the ANOVA analyses. **(A)** For each HPC- and NCL-neuron, a three-way ANOVA was conducted. The leftmost panel depicts the distribution of the explained variances for each factor (Main effects: Phase, Stimulus, Periods) and their interactions (Stimulus*Period, Phase*Period, Stimulus*Phase, 3-way (Phase*Periods*Stimulus)) after a three-way ANOVA for an example cell. The middle panel illustrates how the amount of information elicited by each factor and their interaction is quantified with omega-squared as a proxy. This conversion enabled us to disambiguate the contribution of each factor on the neural responses and accounted once more for the differences between individual neurons e.g., differences in firing rates. Following this procedure, the HPC and the NCL were compared on a population level using a 2-way mixed ANOVA (rightmost panel). **(B)** Description of the amount of information carried by each factor and their interactions for both datasets. HPC-neurons carried more information (∼10 %) than those of the NCL (∼7 %) regarding the different experimental phases (acquisition, extinction, and renewal test). On the other hand, the interaction Stimulus*Phase as well as the 3-way interactions accounted for more information in the NCL than in the HPC. Error bars represent the standard errors of the means. Asterisks mark the statistical significance of post hoc comparisons between the regions using the Tukey HSD test (*p* < 0.05).

Following each ANOVA, we quantified the information carried by the three main factors as well as their interactions based on their effect sizes (omega-squares; see methods section). Using this measure allows us to quantify and compare different neurons with each other with respect to the type of information they encoded. Figure 4B depicts the mean (±SEM) amount of information carried by each factor and interaction for the HPC and the NCL.

With comparable values that indicate the information carried by each factor and interaction for both brain regions, we can now assess whether HPC- and NCL-cells differed with respect to the type of information they carry. For this purpose, we performed a two-way mixed ANOVA to assess the effect of the region (HPC vs. NCL) and the type of information (seven in total: experimental phase, stimulus, periods within a trial, and all their interactions) on the amount of information carried. Although no significant effect of region could be detected (*F*(1,226) = 1.76, *p* = 0.186), we found that the type of information in the experiments did have a significant effect (*F*(6,1356) = 70.3, *p* < 0.001; Figure 4B; Table S1). In addition to the main effects, the interaction between the types of information and the regions was also significant (*F*(6,1356) = 3.44, *p* < 0.01). In summary, three major findings resulted from our initial analysis: (1) The factor phase carried significantly more information in the HPC as compared to the NCL (Tukey test, *p* = 0.046; Table S2). (2) The information accounted for by the interaction Stimulus * Phase was significantly higher in the NCL than in the HPC (Tukey test, *p* = 0.032) (3) The information accounted for by the 3-way interactions was significantly higher in the NCL than in the HPC (Tukey test, *p* < 0.001). In the subsequent sections we will separately elaborate on each of these findings.

### HPC activity shows stronger coding for experimental phases

The amount of information carried by the factor phase was significantly higher in the HPC than in the NCL. Post-hoc comparisons showed that this explained ∼10 % of the total information in the HPC and ∼7 % of the NCL (Tukey test, *p* < 0.001; Figure 4). To ensure that this finding is not related to fading signal quality during our recording session, we controlled for the stability of our recordings. Since no fluctuations in the spike amplitudes could be detected throughout the experiments (Friedman’s test, *χ*^2^(2) = 1.73, *p* = 0.42), we concluded that the changes in the firing rate of these responsive neurons were solely based on the phase of the experiment. Figure 5 exemplifies a cell that carries high information regarding phase (raw data). One important aspect has to be highlighted here: The change in the firing rates cannot be attributed to the mere change in the context. As can be seen in figure 5, the firing rates are elevated also in the ITI, a time window where the context lights are turned off. This means that the experimental phase is represented in the cellular response (cf. figure S6).

**Figure 5.**
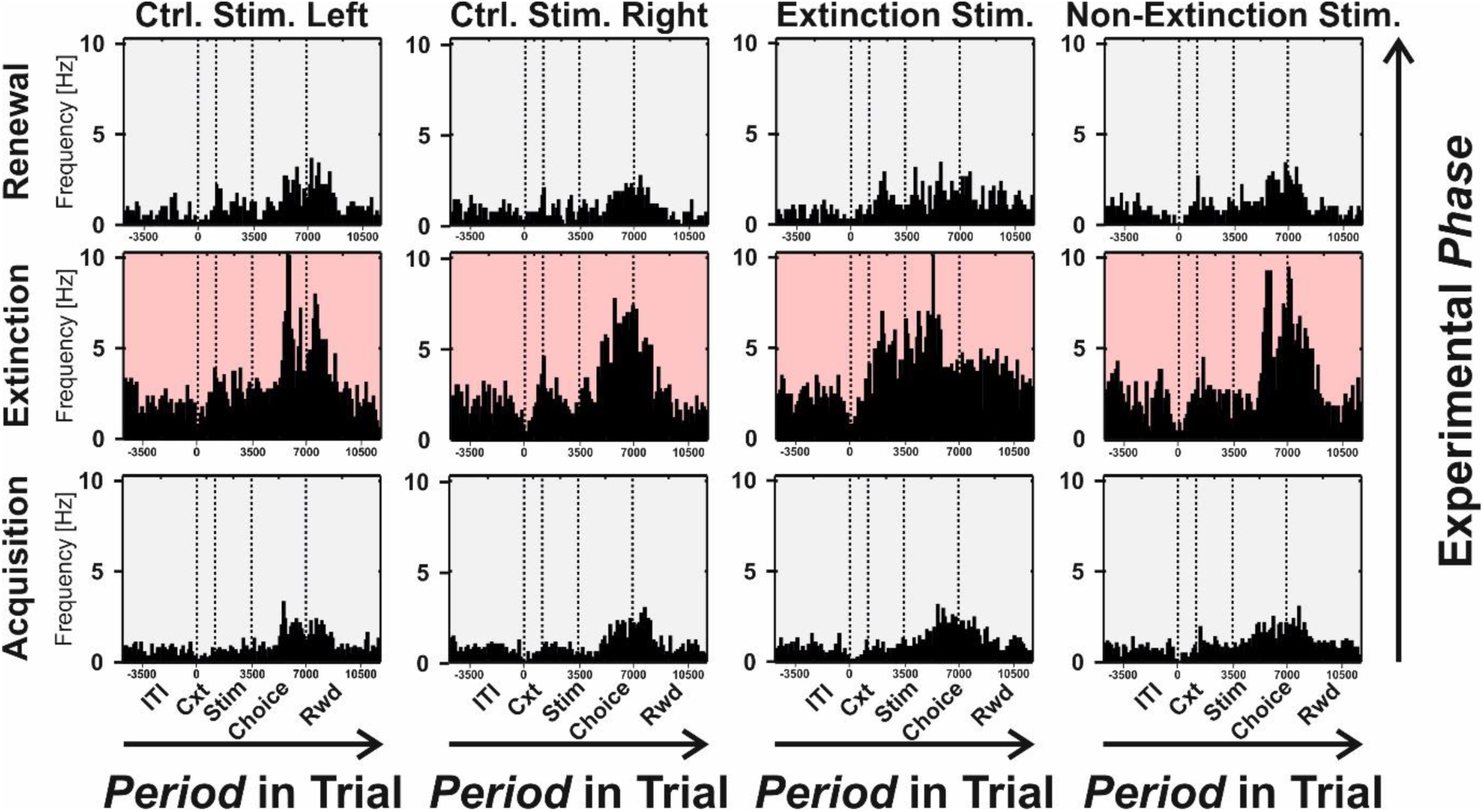
Single unit example from the HPC. The three rows correspond to the acquisition, extinction, and the renewal test phases. The columns represent the individual responses for the different stimuli in the experiment. Each panel provides a peri-stimulus time histogram of the raw data with a bin size of 200 ms. The cell exhibits a strong main effect of phase. More specifically, it increases its FR as the animal enters the extinction phase and decreases its FR again once the animal reaches the renewal phase. Importantly, the changed light condition does not account for this response, since the context light was turned off during the ITI. However, the firing rate remained elevated also during this trial period and throughout the extinction phase (see also figure S6).

### Successive exposure to extinction learning fosters meta-learning

Donoso et al., (2021) investigated the time-course of the behavioral changes throughout the repeated exposure to the context-dependent extinction learning. One of their key findings was that renewal responses decay over successive sessions. This was also the case in our experiments. One potential explanation why the animals started to show reduced renewal responses over time might be a meta-learning process in which the animals slowly start to expect that one of the novel stimuli will be extinguished. To test this idea, we conducted linear mixed models with phase information as dependent variable for each region with pigeons as random factor and days as fixed factor (Figure 6). Although HPC accounted for much of phase information coding (Figure 4), the activity of hippocampal neurons did not change over longer time periods (HPC: *R*^2^ = 0.1, β_phase_ = -0.0004, t(96) = -0.18, *p* = 0.857). By contrast, the amount of phase information in the NCL increased as the animals were tested on consecutive days (NCL: *R*^2^ = 0.07, β_phase_ = 0.004, t(134) = 3.33, *p* < 0.001). We repeated the same analyses for the remaining types of information for both regions and found no significant changes. Therefore, our results suggest that NCL-neurons were able to track the deeper structure of our study, in which always one of the new stimuli underwent extinction and subsequent renewal with concomitant context changes. Most importantly, this meta-learning was processed by NCL- and not by HPC-neurons. To ensure that this effect was not driven solely by the prominent outliers on day 15, we excluded these datapoints and repeated our analysis. Still, we found an increasing amount of phase information in the NCL highlighting the robustness of the found effect (*R*^2^ = 0.068, β_phase_ = 0.003, t(132) = 2.99, *p* = 0.002).

**Figure 6.**
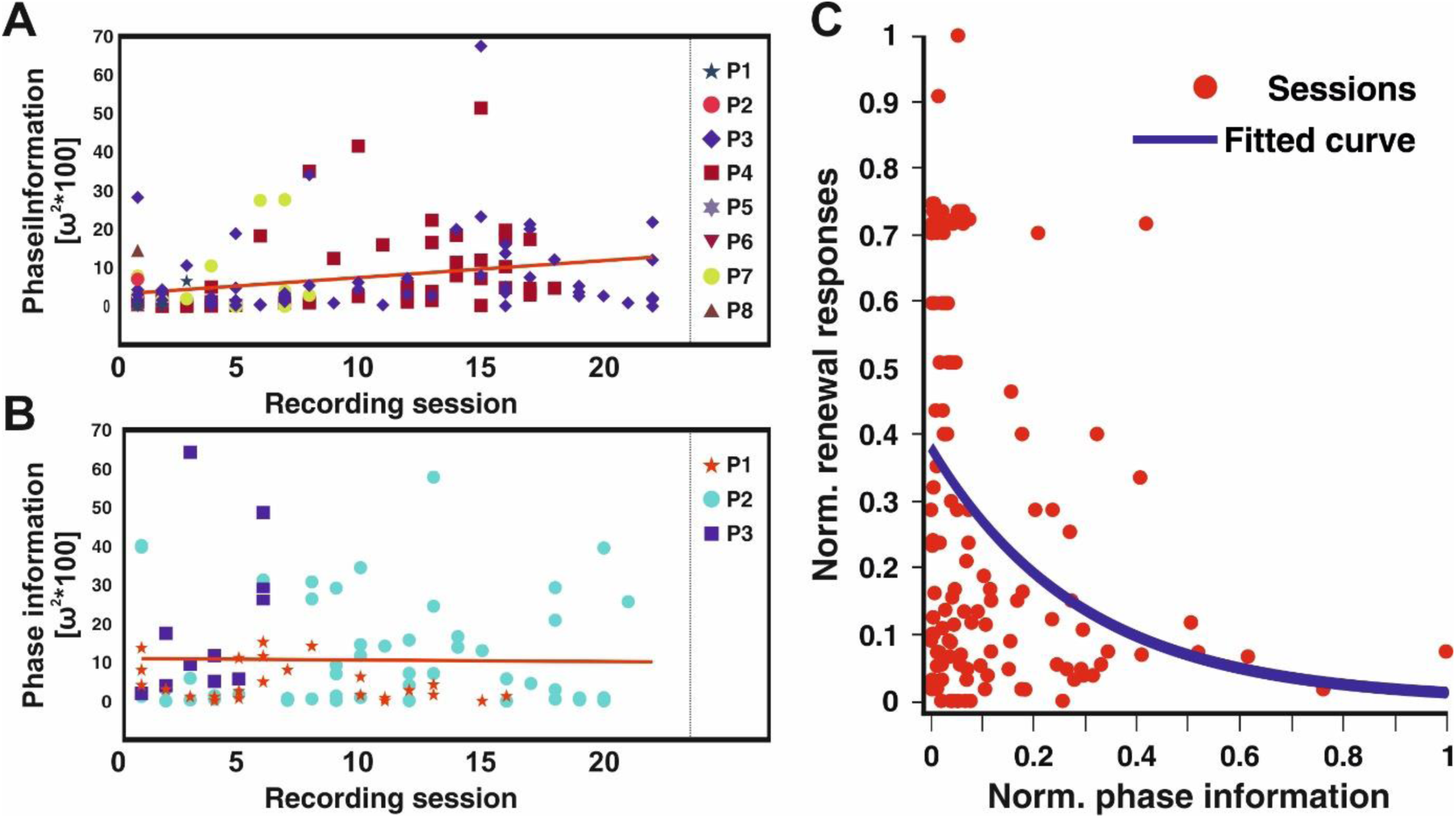
Depiction of the changes in the amount of phase information in both areas in consecutive sessions. **(A)** NCL. **(B)** HPC. The dots correspond to recorded neurons from the respective area and are sorted chronologically. Each pigeon is represented by a different symbol-color combination. The red lines correspond to the fitted regression for each region. While there was no change in the amount of phase information in the HPC over days, the NCL reflected an increase in the amount of phase information it carried. **(C)** The relation between the renewal responses and the phase information in the NCL. Both measures are normalized to the maximum value within each set. We then mapped the renewal responses to the phase information measured in that session. The slope estimate deriving from the linear mixed model regression indicates that the relation between the two measures is disproportional (β_phase_= -23.03): The more phase information there is, the less likely the renewal responses become. We found that the best model that fits our data is exponential and is depicted here with the blue curve.

### The amount of renewal responses inversely correlates with phase information

As shown above, NCL neurons signal more phase information over time. Consequently, we now analyzed if an increase of HPC and NCL responses is related to the number of responses during the renewal phase using separate linear mixed model regression for each session as a dependent variable. Thereby, phase information and the information carried by 3-way interactions served as predictors. Due to the nested structure of our experiment, we introduced pigeons and sessions as random effects in a hierarchical manner. We found that no type of information had predictive power on the renewal responses in the HPC (*R*^2^ = 0.31, β_phase_ = 4.15, t(95) = 0.86, *p* = 0.37; β_3-way_ = -148.55, t(95) = -1.44, *p* = 0.15), however, the phase information and the information regarding 3-way interactions were significant predictors of the renewal responses in the NCL (*R*^2^ = 0.49, β_phase_ = -24.17, t(133) = -2.81, *p* < 0.01; β_3-way_ = -119.41, t(133) = -1.96, *p* = 0.05). To investigate the contributions of each single factor, we dropped the 3-way interactions as a second predictor and ran another model. We found that this reduced model (*R*^2^ = 0.48, β_phase_ = -23.03, t(134) = -2.64, *p* < 0.01) was on par with the former one (|Δ*R*^2^| = 0.01). This result indicates that sessions with high neuronal phase modulation predict fewer renewal responses and vice versa (Figure 6C).

### NCL integrates ascending projections

As a second result, we found that specific combinations of main factors of the ANOVA were effective in driving the cells’ activation profiles. Many neurons responded to more than one task parameter and showed mixed selectivity. This type of coding property is believed to be highly flexible, maximizes computational power, and allows an easy readout of information by downstream neural circuits (Tye et al., 2024). Figure 7 illustrates a cell with a strong 3-way interaction from the NCL dataset. This cell was specifically active during the extinction phase of the experiment and responded to only one stimulus during the phase of the reward delivery resulting in a 3-way interaction (Phase*Stimulus*Period). We found that the amount of these three-way interactions was higher in the NCL than in the HPC (Tukey test, *p* < 0.001; Figure 4).

**Figure 7.**
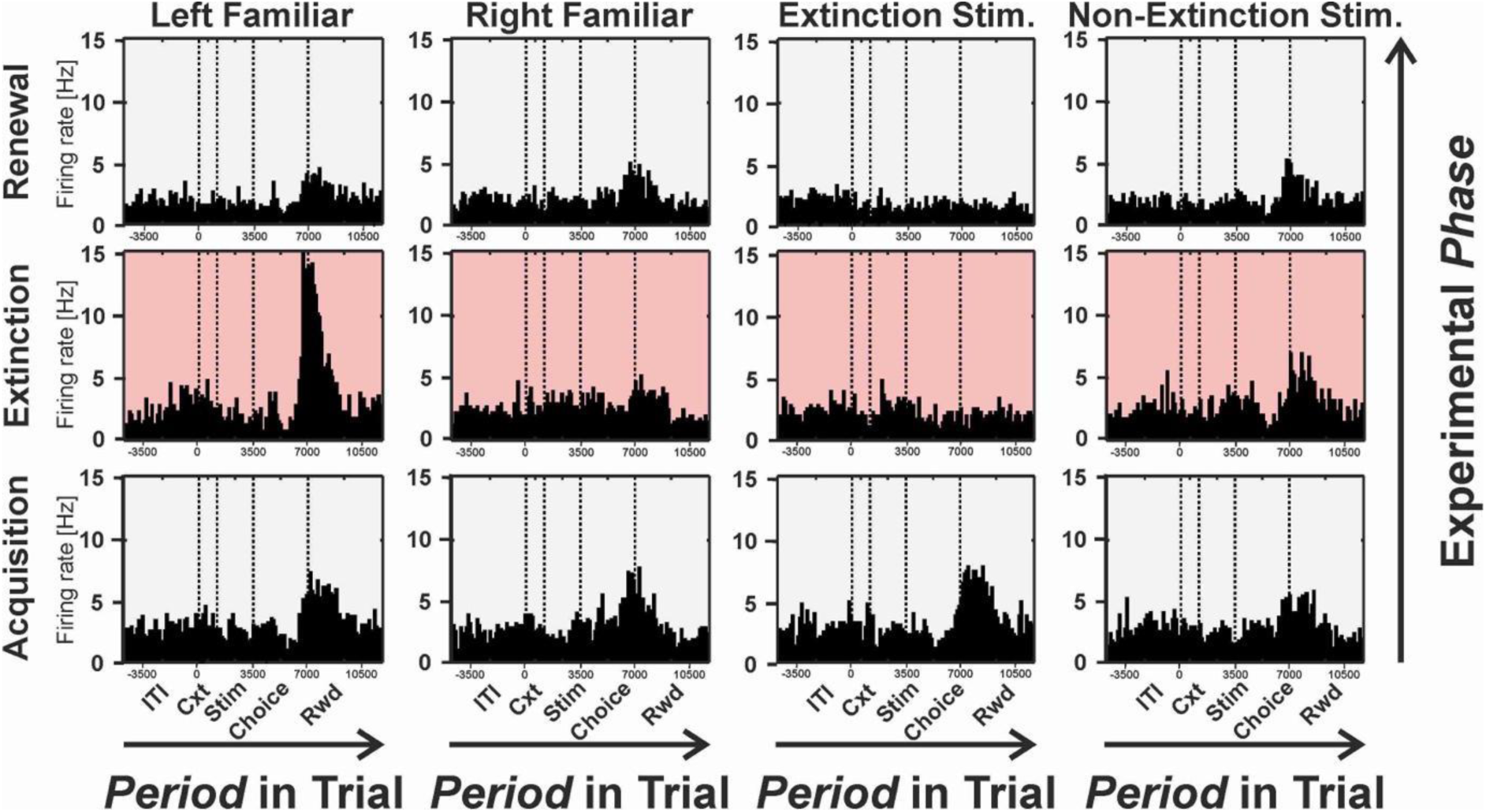
Single unit example from the NCL. As in figure 7, the rows correspond to the acquisition, extinction, and the renewal phases. The columns represent the individual responses for the different stimuli in the experiment. Each panel provides a peri-stimulus time histogram of the raw data with a bin size of 200 ms. The cell has a very specific time window where it becomes responsive. It increases its FR in response to one of the control stimuli after the reward delivery only in the extinction phase, illustrating the increased specialization of the NCL neurons.

Yet, up to now, our analysis does not allow to disentangle which factors were mixed and altered the cells’ responses at a single trial resolution. To clarify this, we computed multiple linear regressions with 13 distinct task parameters as predictors (cf. Figure 1E and see section *Neural data analysis*) and raw spike counts as dependent variables. Since most of the sampled cells upregulated in their FRs (cf. Figure 3), we focused only on the positive β-values in the remaining analysis. The results including the negative β-values can be found in the supplementary figure S6.

For both regions, mixed responses to several task parameters were not random. Instead, we found that HPC and NCL show remarkably distinct patterns of mixed selectivity of different parameters in our experiments (Figure 8): HPC cells formed two functional clusters mainly implicated in the representation of upcoming decisions and the representation of context and stimuli. No significant co-modulation of outcome parameters was found in the HPC. Responses for different contexts and stimuli were found to be densely clustered together in the HPC. These results in conjunction with our findings from the previous section suggest that the information about contexts and stimuli as present in the hippocampus cannot yet be converted into a binary decision explaining the inverse correlation of phase information and renewal responses.

**Figure 8.**
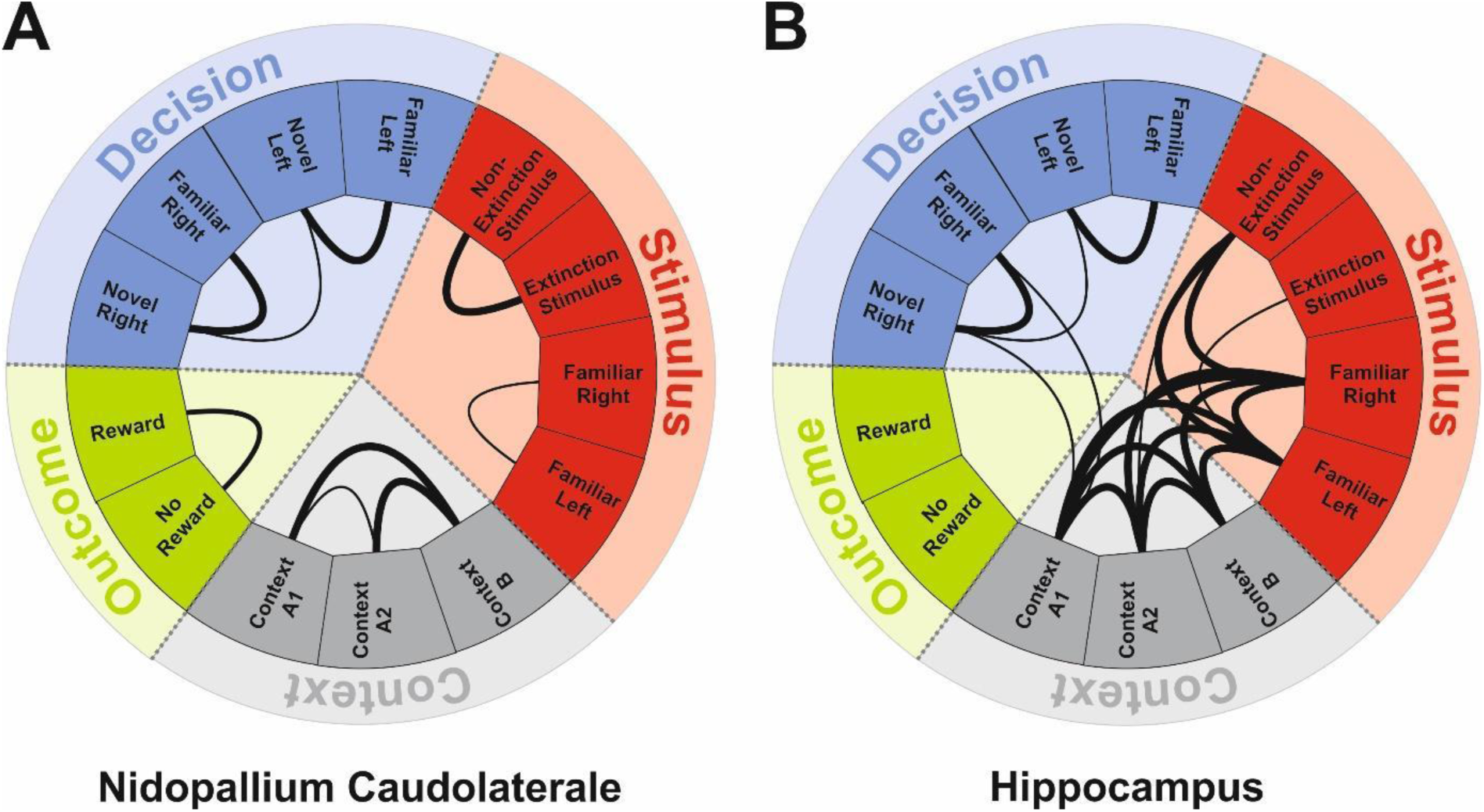
Connectograms depicting significant co-responses of neurons. To explore which parameters coupled and modulated the cells’ activity, we regressed the FRs with the task parameters we employed in the experiment (cf. Figure 1E). Thick or thin connections between each parameter indicate statistical significance on the population level (p = 0.01, and p = 0.001, respectively). In the NCL, significant connections built functional ensembles that corresponded to the different experimental parameters. In the HPC there is no significant co-response of the outcome phase. Further, co-responses of context and stimuli overlap.

In contrast, mixed selective neurons of the NCL formed four significant functional clusters that correspond to the task parameters at hand. This organization of cell responsiveness in the NCL distinguishes between contexts, stimuli, decisions, and outcomes. Especially within these task parameters, co-responses of neurons can be interpreted as decision bifurcations (Figure 8). This response profile is consistent with the proposed role of the NCL as an associative brain region with high levels of abstraction (Güntürkün et al., 2024).

The amount of structure within the connectograms can be quantified using modularity coefficients (Newman, 2006). For the NCL, the dense coupling within task parameters void of connections between other parameters is reflected in a high modularity coefficient (µ^NCL^ = 0.46). In contrast, co-responses in the HPC appear to have less specificity resulting in a lower modularity coefficient (µ^HPC^ = 0.11). Our findings suggest that mixed selective information about stimuli and contexts is present and provided by the HPC. This information is integrated by the NCL to guide adaptive behavioral decisions.

## Discussion

In the present study, we investigated the neural correlates of contextual cue integration and decision dynamics into the avian extinction network. To this end, pigeons were trained in an appetitive extinction learning paradigm (Packheiser et al., 2019), while we recorded single-unit responses from the HPC and the NCL. Although our recordings from both areas did not take place simultaneously, we observed nearly identical behavioral performances in both datasets, and thus compared the neural correlates of extinction learning in the pigeon HPC and NCL. We discovered three key findings. First, context was represented by HPC neurons based on the representation of experimental phases. Second, NCL neurons represented a meta-learning effect with which the animals slowly started to exploit the deeper structure of our task to reduce renewal responses. Third, HPC and NCL cells displayed different combinations of mixed selectivity responses, that possibly aided stimulus integration and decision making, respectively. We will discuss these points, one by one.

### Hippocampal neurons represent experimental phases based on context

The first key result of our study is that population-level HPC neurons represent the successive experimental phases stronger than the NCL. This effect, however, only emerged after contextual cues started to become relevant during extinction (cf. Supplementary figure S3). To understand the relevance of this finding, it is important to recall the function of context in extinction learning: In the moment extinction starts, the meaning of the S+ becomes ambiguous since it ceases to predict reward but instead starts to predict its absence. According to the Attentional Theory of Context Processing, organisms tend to ignore context information as long as they have no predictive power (Rosas & Callejas-Aguilera, 2006; Ogallar et al., 2017). However, when the CS+ loses its predictive properties during extinction, a search for better predictors starts and attention is shifted to the context. Since the context change coincides with the change of reward properties, context-specific processing starts during the phase of extinction (context B). Importantly, the physical properties of the context cannot account for this type of response. Even within the ITI where the context signal is turned off, the phase information is maintained. Thus, it is the experimental phase that is signaled in the cellular responses (cf. figure 7).

Studies in mammals have demonstrated that especially HPC neurons reflect contextual information about the physical environment (Plitt & Giocomo, 2021; Zhao et al., 2020). Our results show that the same holds for pigeons (Figure 3). Thus, based on contextual changes, the HPC provides the extinction network with temporal information about the sequences of events during extinction. Most importantly, avian HPC neurons incorporate contextual cues into the extinction network on a temporal scale, i.e. across different phases of the experimental paradigm. Thus, the representation of sequential events possibly serves to integrate contextual changes into the extinction learning process - a pattern that is consistent with findings in the mammalian hippocampus (Sun et al., 2020; Bulkin et al., 2020; Rubin et al., 2015). These response profiles mostly emerged during extinction and renewal phases since only then context changes coincided with expectancy violations (Packheiser et al., 2021). Our findings align with behavioral studies in pigeons that highlighted the increased relevance of contextual cues when predictions regarding stimulus-outcome associations break down (Starosta et al., 2014). Thus, when faced with uncertainty, contextual information gains prominence, aiding in the adjustment of learned responses (Bernal-Gamboa et al., 2014; Rosas & Callejas-Aguilera, 2006).

### NCL neurons represent meta-learning based on a long-term integration of experimental phases

Part of our second key finding is that the total number of renewal responses decreased across subsequent sessions (cf. Supplementary figures S7 & S8). Thus, based on their daily experiments, our pigeons learned to expect that one of the acquired novel stimuli was going to be no longer rewarded when context changes. This implies that the animals acquired a deeper rule of our experimental design – a process called ‘learning-to-learn’ or meta-learning. The fundamental aspect of meta-learning is its long-term experience dependency that allows for inductive biases or knowledge that speeds up future learning (Wang, 2021). To our knowledge it was first demonstrated by Harlow (1949) in non-human primates who received two new objects in every block of six trials. Since only one was rewarded, the optimal strategy would be one-shot learning in which the monkey randomly selects one of the objects and then proceeds based on the reward outcome of this first trial. Harlow’s monkeys demonstrated meta-learning but required ca. 300 blocks for a close-to-optimal strategy. Hattori et al. (2023) could show that also mice can slowly learn to acquire the deeper structure of a serial reversal learning task by gradually shaping the population value coding properties of orbitofrontal neurons to guide the ongoing behavioral policy. Various studies make it likely that the dopaminergic input of the prefrontal cortex plays a key role for meta-learning. For instance, in monkeys that learn a reversal task, dopaminergic neurons signal both the just experienced value and in addition the inferred value of the non-chosen stimulus, thereby allowing to process alternative decisions (Bromberg-Martin et al., 2010). Consequently, pharmacological enhancing catecholamine functions in human participants increases their meta-learning rate (Cook et al., 2019).

Here, we found that NCL neurons carried meta-learning related information about the experimental phases when sessions progressed (Figure 6A). This neural finding was accompanied by the behavioral decline of renewal responses in exactly those late sessions. Like in mammals, the NCL is densely innervated by dopaminergic fibers (von Eugen et al., 2020) that modify the activity patterns of excitatory principal neurons (Durstewitz et al., 1998; Kröner et al., 2002), thereby learning-dependently altering dopamine receptor expression profiles and cognitive properties (Herold et al., 2008; 2012). It is important to note that experimental phase information is present in the HPC from early sessions on and drives contextual changes of extinction behavior without varying over months of testing. The same information, however, drives meta-learning in NCL. This means that the same information is processed by HPC and NCL in different ways to enable complementary cognitive processes.

The finding that the phase information predicts renewal responses in NCL but not in HPC contrasts with findings in mammals that show that neural activities in both areas are associated with the emergence of renewal following extinction learning (Milad & Quirk, 2002; Lacagnina et al., 2019; Lissek et al., 2016). This highlights functional dissimilarities between birds and mammals (Zhao et al., 2020; Kelemen & Fenton., 2010; Wills et al., 2005; Herold et al., 2014). It is yet unclear what drives this difference. One possibility is the absence of a direct anatomical connection between HPC and NCL (Shanahan et al., 2013). Instead, their interaction could be mediated via polysynaptic projections of different brain areas. In contrast, the mammalian HPC is connected to diverse subareas of the prefrontal cortex (Lavanex et al., 2002; Rosene & Van Hoesen, 1977) and these projections play a crucial role in the transmission and integration of hippocampal representations within the PFC (Spellman et al., 2015; Tamura et al., 2017; McClelland et al., 1995; Hattori et al., 2023; Hunt et al., 2018; Sul et al., 2010; Miller & Cohen, 2001; Samborska et al., 2022). Differences might possibly also be rooted in intrahippocampal connectivity patterns (Atoji & Wild, 2004; Rook et al., 2023). But despite these differences to mammals, birds seem to have implemented complementary computations in HPC and NCL that exploit the information of experimental phases to achieve contextual integration of extinction learning and meta-learning, respectively.

### Different mixed selectivity patterns in hippocampus and NCL

Our third key finding were the different coding schemes of HPC and NCL. Although both regions showed mixed selectivity, we detected important differences. HPC cells formed only two functional clusters: one represented relevant stimuli, be they contextual or cue-related, while the other coded for upcoming decisions. Similarly, also the mammalian HPC plays a key role in stimulus-stimulus associations, both within a sensory system (Billig et al., 2022), as well as when connecting cues across sensory domains (Borders et al., 2017; Quiroga et al., 2005). Most importantly, these cues are bound into mnemonic representations across time gaps and space (Staresina & Davachi, 2009). Since activity patterns of a subfraction of hippocampal place cells gradually change over daytime, the hippocampus can bind sensory cues across contexts along a mental timeline (Rubin et al., 2015). In the mammalian HPC, these processes could provide the computational means to distinctly code for the change of reward-related properties of stimuli during acquisition and extinction and so contribute to the integration of contextual cues into the extinction network (Lacagnina et al., 2019; Maren et al., 2013; Phillips & LeDoux, 1992; Zhao et al., 2020). Our finding that the avian HPC can integrate all stimuli (contextual, cue-related) by mixed selectivity patterns, could serve a similar computational means to subserve the tight stimulus-stimulus associations that are needed for contextual coding.

Besides the hippocampal contribution, the involvement of the PFC in extinction learning is well established. It is generally assumed that the PFC uses task information to adaptively initiate behavioral decisions, particularly in situations of ambiguity or conflict (Gonzalez & Fanselow 2020). These flexible properties of the PFC are thought to be realized by neurons that integrate and evaluate all relevant task variables by mixed selectivity (Rigotti et al. 2013). In contrast to mixed selectivity of HPC neurons, those of the NCL formed four significant functional clusters that correspond to the task parameters contexts, cues, decisions, and outcomes. This kind of differential representation should allow flexible neural representations with which context information can easily modify the value of cues during extinction (Tye et al., 2024). This would agree with the proposed role of the NCL as an associative brain region with high levels of abstraction that are used to weigh and decide between response alternatives (Güntürkün et al., 2021; 2024).

## Summary

In conclusion, our data provides evidence how contextual cues are incorporated into the avian extinction network, giving rise to context-dependent renewal of the previously extinguished behavior and long-term meta-learning of the deeper structure of our task. Hereby, the HPC integrates context and cue information that is subsequently exploited by the executive forebrain area NCL to guide adaptive decision-making during all phases of extinction learning. In both regions, mixed selective cells were found that differed in their response properties. This feature enabled the complementary computational properties of hippocampus and ‘prefrontal’ NCL to divide and conquer the task.

## Supporting information

All supplemental materials

## Acknowledgments

This work was supported by the Deutsche Forschungsgemeinschaft (DFG, German Research Foundation) through SFB 1280 (A01, A14, A19 and F01) project number 316803389.

