## Supplementary material for "Divide and conquer: How avian “prefrontal” and hippocampal neurons process extinction learning in complementary ways": All supplemental materials: Sevincik_Suppl.pdf

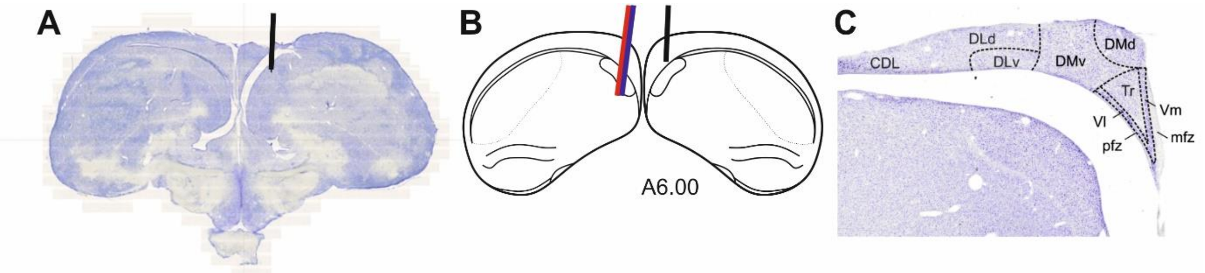


**S1. (A)** Example slice illustrating the electrode track. (B) Individual track reconstruction. (C) Subdivisions of the hippocampus.

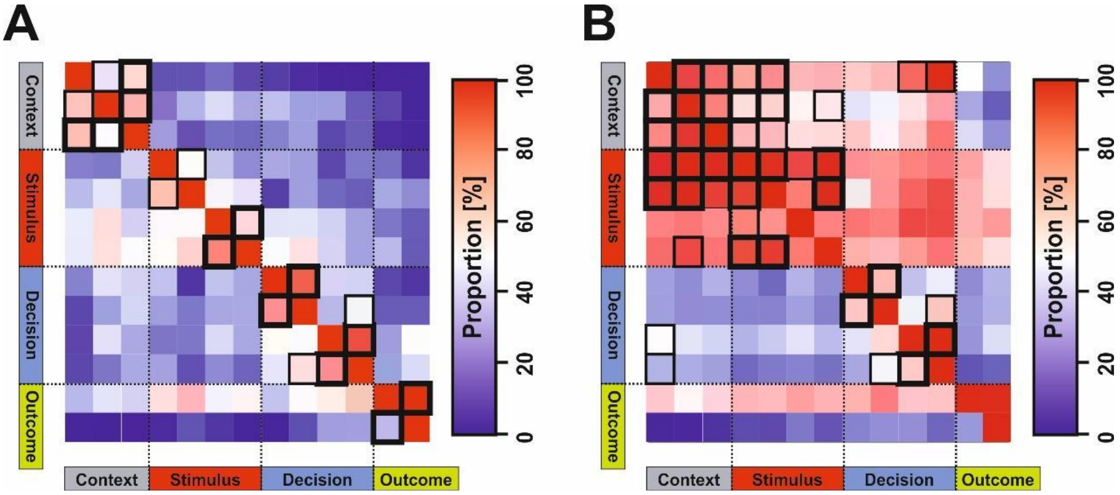


**S2**. Proportion of co-responsive cells for each pair of predictors in the NCL (A) and HPC (B). Proportions correspond to the number of neurons responding to the predictor specified by the row, divided by the total number of cells responding to the predictor specified by the column. Black squares indicate significant co-responses (thin squares, p = 0.01; thick squares, p = 0.001).

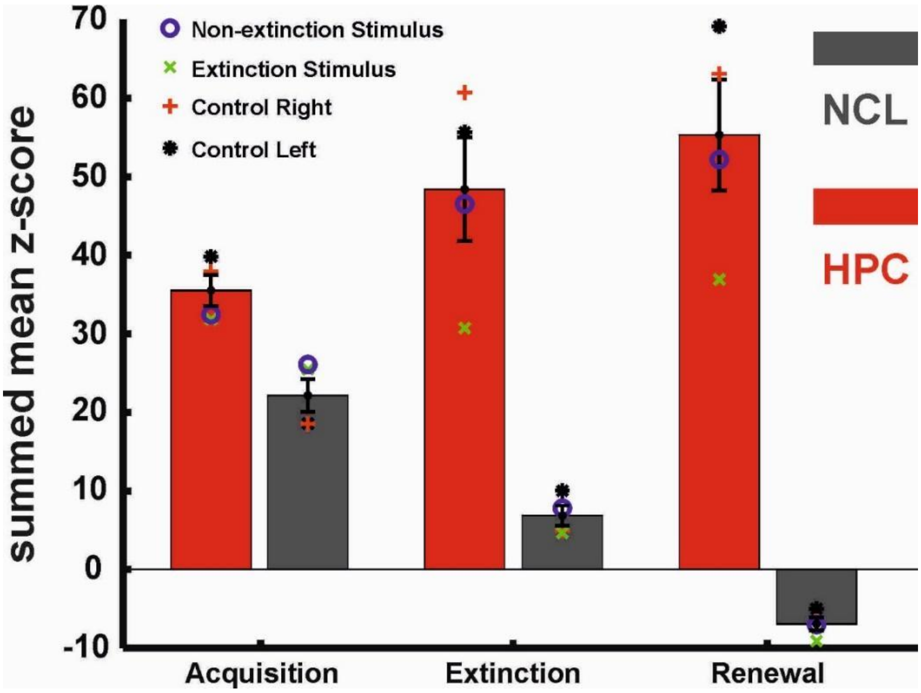

**S3.** Depiction of the overall neural responses for all stimuli during the different phases of the experiment for the HPC and the NCL. The responses in the HPC were generally stronger as compared to the NCL response profiles. While the response strength increased throughout the experimental phases in the HPC, it declined in the NCL.

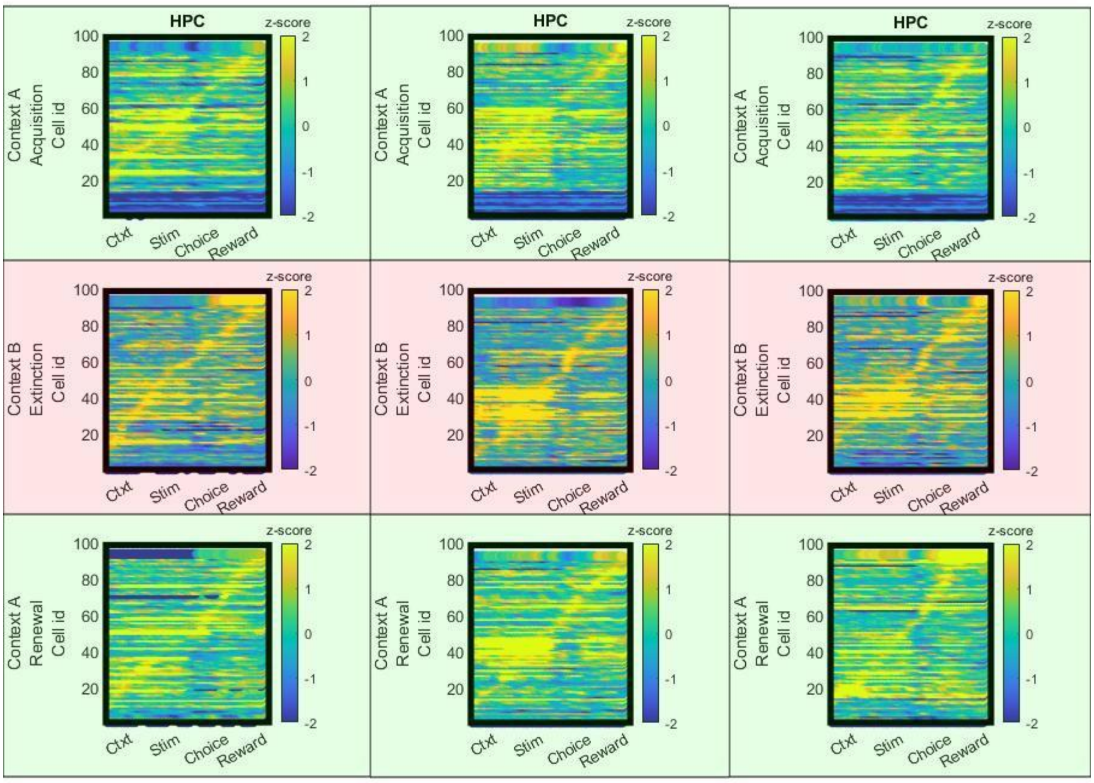


**S4**. Population responses of the HPC neurons for the control left, control right, and novel non-extinction stimuli respectively.

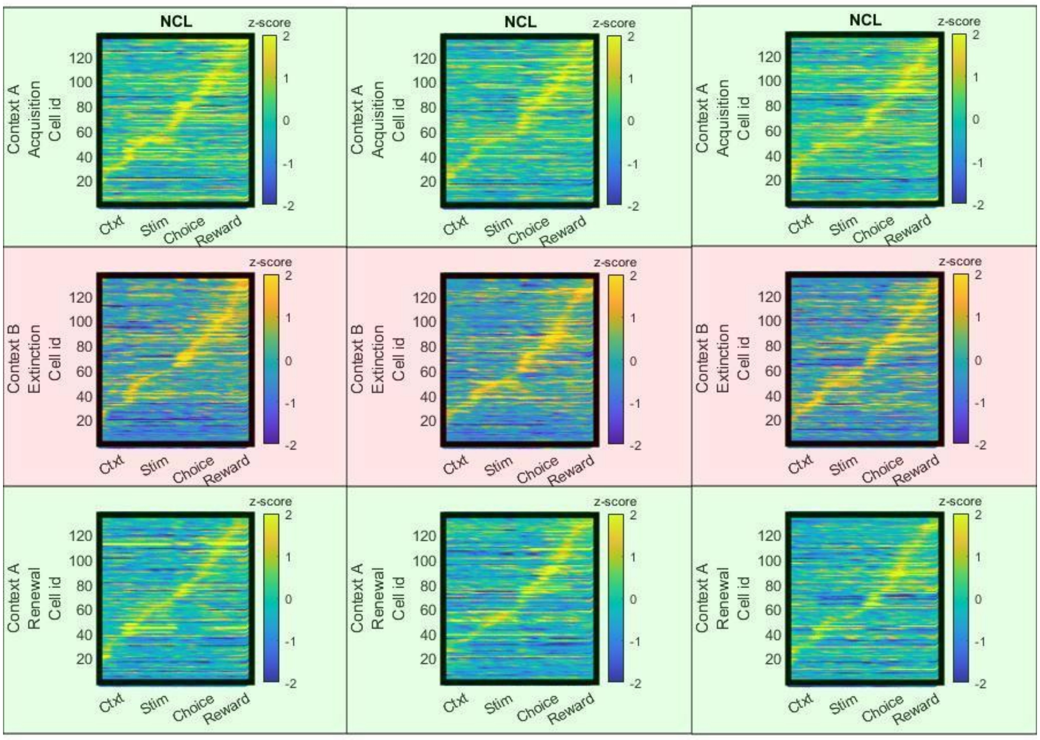


**S5**. Population responses of the NCL neurons for the control left, control right, and novel non-extinction stimuli respectively.

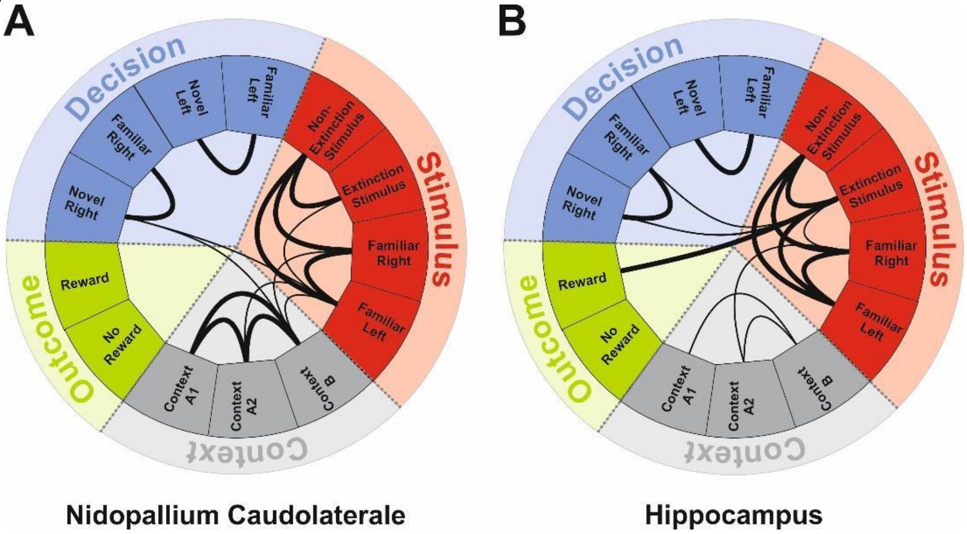


**S6**. Connectivity analysis of negative activations. (A) NCL. (B) Hippocampus.

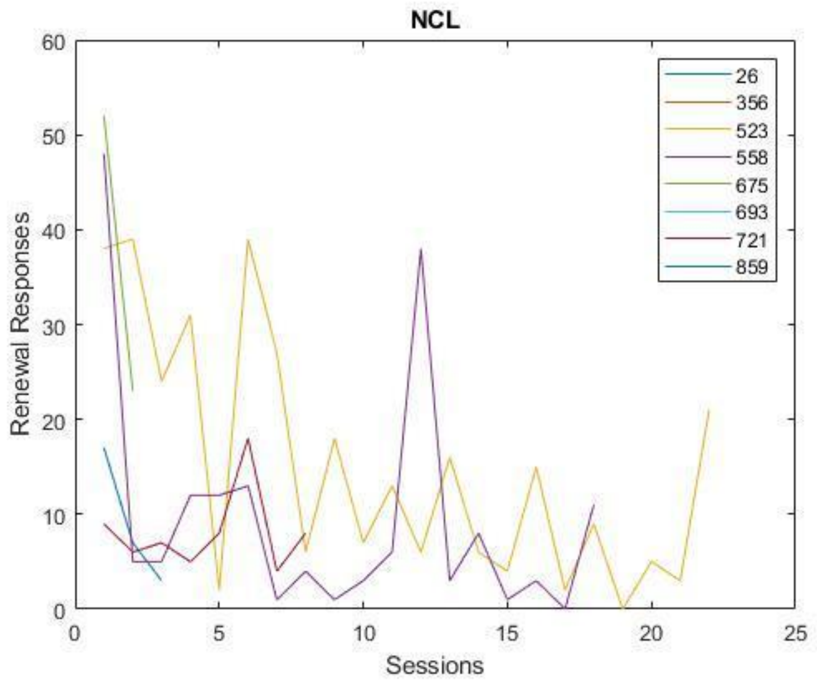

**S7**. Renewal responses in the NCL as a function of session repetition.

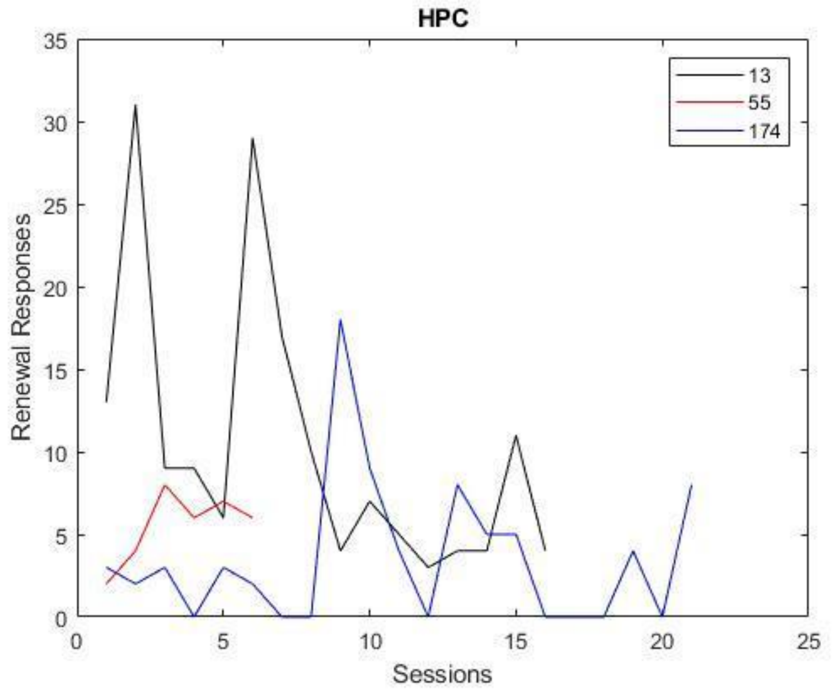

**S8**. Renewal responses in the HPC as a function of session repetition.

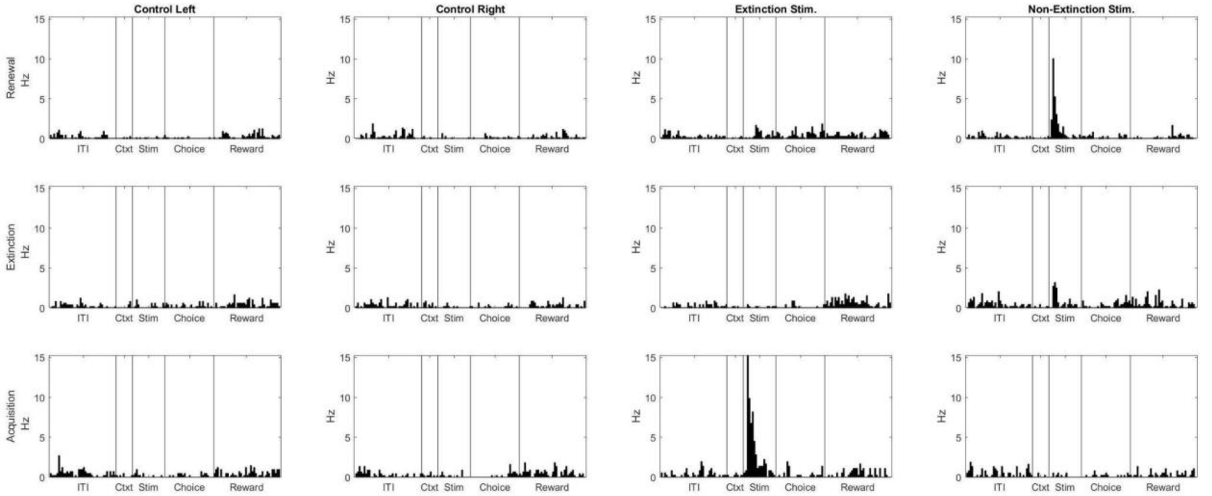


**S5**. A second example cell displaying a relatively strong 3-way interaction from the NCL dataset. The bin size was set to 100 ms.

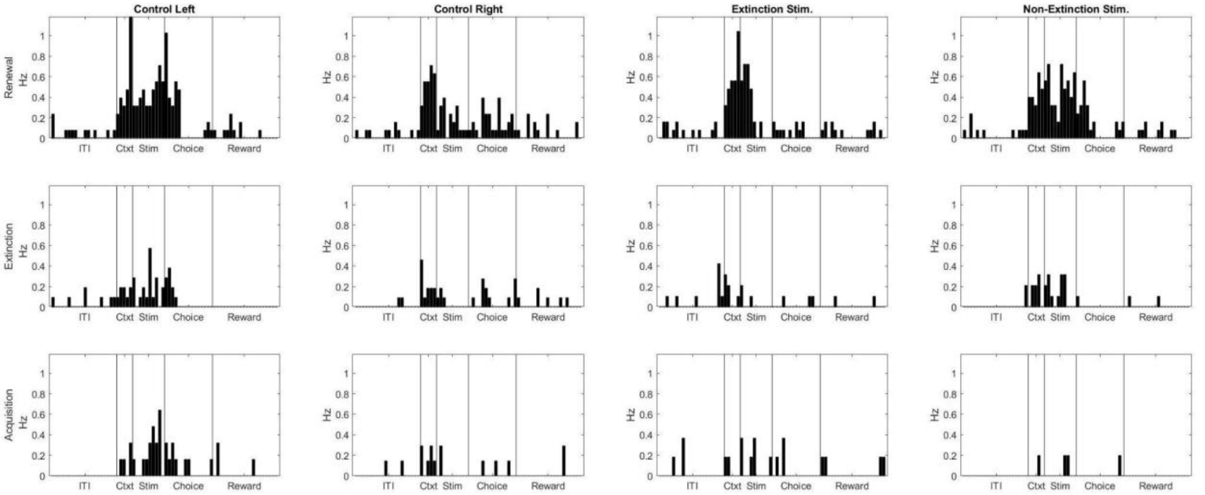


**S6**. A second example cell reflecting a relatively strong phase information from the HPC dataset. The bin size was set to 100 ms.

### Table S1

*Two-way Mixed ANOVA Results*

**Between-subject effects**

| Source | Sum of Squares | df | Mean Square | F | Sig. |
|---|---|---|---|---|---|
| Intercept | 1.039 | 1 | 1.039 | 354.040 | <.001 |
| Region | 0.005 | 1 | 0.005 | 1.757 | 0.186 |
| Region * Information Type | 0.055 | 6 | 9 | 3.440 | 0.002 |
| Error (Region) | 0.663 | 226 | | | |

**Within-subject effects**

| Source | Sum of Squares | df | Mean Square | F | Sig. |
|---|---|---|---|---|---|
| Information Type | 1.115 | 6 | 0.186 | 70.293 | <.001 |
| Error (Information Type) | 3.585 | 1,356 | | | |

**Table S2**
*Results of Multiple Post hoc Tukey Tests*

| Information Type | NCL | HPC | Difference | StdErr | pValue | Lower | Upper |
|---|---|---|---|---|---|---|---|
| Phase | 0 | 1 | -0.0321 | 0.0161 | 0.0460 | -0.0636 | -5.6841e-04 |
| Phase | 1 | 0 | 0.0321 | 0.0161 | 0.0460 | 5.6841e-04 | 0.0636 |
| Stimulus | 0 | 1 | -2.8919e-04 | 0.0026 | 0.9109 | -0.0054 | 0.0048 |
| Stimulus | 1 | 0 | 2.8919e-04 | 0.0026 | 0.9109 | -0.0048 | 0.0054 |
| Period | 0 | 1 | -0.0043 | 0.0079 | 0.5885 | -0.0197 | 0.0112 |
| Period | 1 | 0 | 0.0043 | 0.0079 | 0.5885 | -0.0112 | 0.0197 |
| Stimulus * Phase | 0 | 1 | 0.0020 | 9.2570e-04 | 0.0324 | 1.6638e-04 | 0.0038 |
| Stimulus * Phase | 1 | 0 | -0.0020 | 9.2570e-04 | 0.0324 | -0.0038 | -1.6638e-04 |
| Stimulus * Period | 0 | 1 | 0.0021 | 0.0030 | 0.4836 | -0.0038 | 0.0079 |
| Stimulus * Period | 1 | 0 | -0.0021 | 0.0030 | 0.4836 | -0.0079 | 0.0038 |
| Phase * Period | 0 | 1 | 0.0014 | 0.0016 | 0.3745 | -0.0017 | 0.0046 |
| Phase * Period | 1 | 0 | -0.0014 | 0.0016 | 0.3745 | -0.0046 | 0.0017 |
| 3 way | 0 | 1 | 0.0055 | 0.0016 | 8.2854e-04 | 0.0023 | 0.0087 |
| 3 way | 1 | 0 | -0.0055 | 0.0016 | 8.2854e-04 | -0.0087 | -0.0023 |
